# CD4 T cell contact drives macrophage cell cycle progression and susceptibility to lentiviral transduction

**DOI:** 10.1101/2024.01.15.575666

**Authors:** Petra Mlcochova, Raphael Heilig, Roman Fischer, Ravindra K. Gupta

## Abstract

Macrophages are typically quiescent cells residing in G0, though tissue macrophages have been shown to proliferate locally in tissues; we previously demonstrated that differentiated monocyte derived macrophages (MDM) can be stimulated to re-enter G1 phase of the cell cycle from G0, without cell division. Entry into G1 correlates with an increase in CDK1 expression which phosphorylates the deoxynucleotide-triphosphate hydrolase SAMHD1 at position 592. SAMHD1 not only regulates cellular dNTP levels, but is also a restriction factor for virus replication of HIV-1 and DNA viruses. Here we show that contact with autologous CD4 T cells leads to antigen-independent macrophage cell cycle progression from G0-G1, accompanied by expression of cell cycle associated proteins, including CDK1, and the activation of the canonical MEK-ERK pathway. Further, macrophage cell cycle progression can be blocked not only by anti-cancer drugs targeting the MEK-ERK axis such as Palcociclib, but also by pre-treatment with EGFR antibody, providing additional evidence for cell surface interactions driving proliferative responses. Cell contact with uninfected CD4 T cells renders macrophages ten-fold more susceptible to transduction with VSV-G pseudotyped HIV-1 particles. These experiments demonstrating cell cycle progression following antigen independent T cell contact delineate a new phenotype of macrophage, with distinct distribution of cell surface marker distribution from M0/M1/M2. The deactivation of SAMHD1 antiviral activity following cell contact may underlie observations that lymphoid tissues remain a difficult to treat HIV reservoir even in the face of antiretroviral therapy.

**Significance statement:** Macrophages play critical roles across health and disease and are targets of a range of pathogens, including viruses. SAMHD1 is an antiviral restriction factor that is active when cells in G0. Macrophages typically in the G0 cell cycle state, though macrophages have been observed to proliferate. Here we show that contact with autologous CD4 T cells leads to antigen-independent macrophage cell cycle progression from G0-G1, accompanied by expression of cell cycle associated proteins, including CDK1, and the activation of the canonical MEK-ERK pathway. These experiments demonstrating cell cycle progression following antigen independent T cell contact delineate a new phenotype of macrophage, with distinct distribution of cell surface marker distribution from M0/M1/M2. CDK1 associated SAMHD1 phosphorylation at T592 in macrophages renders them hyper-susceptible to HIV-1 transduction and therefore macrophage-T cell interactions typical of HIV sanctuary sites such as lymph nodes and spleen may promote HIV persistence.

## Introduction

Macrophages are cells implicated in a diverse array of functions such as defence against pathogens, tissue repair /homeostasis, and anti-cancer activity^1,2^. In addition, macrophages play a role as antigen presenting cells for the adaptive immune response. As such macrophages form ‘immunological synapses’ with CD4 T cells during MHC II dependent antigen presentation. These involve contact between the two cells via a number of interacting surface proteins including LFA-1 and ICAM-1 in addition to peptide loaded MHC interaction with the cognate TCR on CD4 T cells. Cell polarisation and reorganisation/protein clustering occurs after continual and repeated contact^3^ in order to facilitate signalling in the T cell^4^ and triggering of an immune response that includes release of IL2^5^. Much less is known regarding antigen independent contact between macrophages and T cells, although they can reciprocally respond to signals from each other to induce inflammatory responses.

Both T cells and macrophages are targets for HIV-1^6^; cell cycle progression renders both cell types susceptible to infection via CDK-mediated phosphorylation of SAMHD1, a cellular deoxynucleotide-triphosphate hydrolase and lentiviral restriction factor^7,8^. Whilst T cells often proliferate in response to stimulation and become highly susceptible to HIV-1, macrophages typically reside in G0 and are rather refractory to HIV-1^6,9^.

We previously demonstrated a laboratory-based method to stimulate cell cycle progression from G0 to G1 phase in monocyte-derived macrophages (MDM) via activation of the canonical Ras-Raf/MEK/ERK mitogen associated pathway, with an increase in CDK1 expression^9^. More recently we showed that low oxygen environment can also drive cell cycle progression via HIF2a activation of the same mitogen associated pathway^10^. By contrast, danger signals such as ^11^and cytotoxic agents^12^ can limit cell cycle progression in macrophages and block lentiviral transduction^9^. Here we investigated physiological means by which macrophages may be activated to enter the cell cycle and become more susceptible to transduction. Given known transmission of HIV from T cells to macrophages^13,14^, we hypothesised that infection of macrophages may be facilitated by contact with CD4 T cells as found in tissue reservoirs of HIV such as lymph nodes^15^ and spleen.

## Results

Monocytes were derived from blood apheresis cones of healthy individuals by adhesion in tissue culture plates before treatment with MCSF to induce differentiation into macrophages for 6 days. Cells were further exposed for 2 days to control conditions (M0), stimulated with LPS/IFNgamma to induce M1 macrophages or IL4/IL-13 to induce M2 polarised macrophages. A fourth group of cells were co-cultured with autologous CD4+ T cells.

Monocyte-derived macrophages were then washed to remove CD4 T cells and either (i) stained for cell surface (polarisation) markers (ii) infected with HIV-1 VSV-G pseudotyped virus (PV), or (iii) stained for markers of cell cycle progression. Cell surface macrophage polarisation markers were investigated under each condition (Figure 1A). As expected M1 polarised macrophages upregulated CD80 and CD86 (Figure 1B); M2 macrophages expressed higher levels of CD86 and CD209 compared to unstimulated macrophages. Whilst macrophages co-cultured with autologous CD4 T cells did not show clear polarisation into either M1 or M2 macrophages, there was upregulation of CD209 similar to that observed in M2 macrophages.

**Figure 1:**
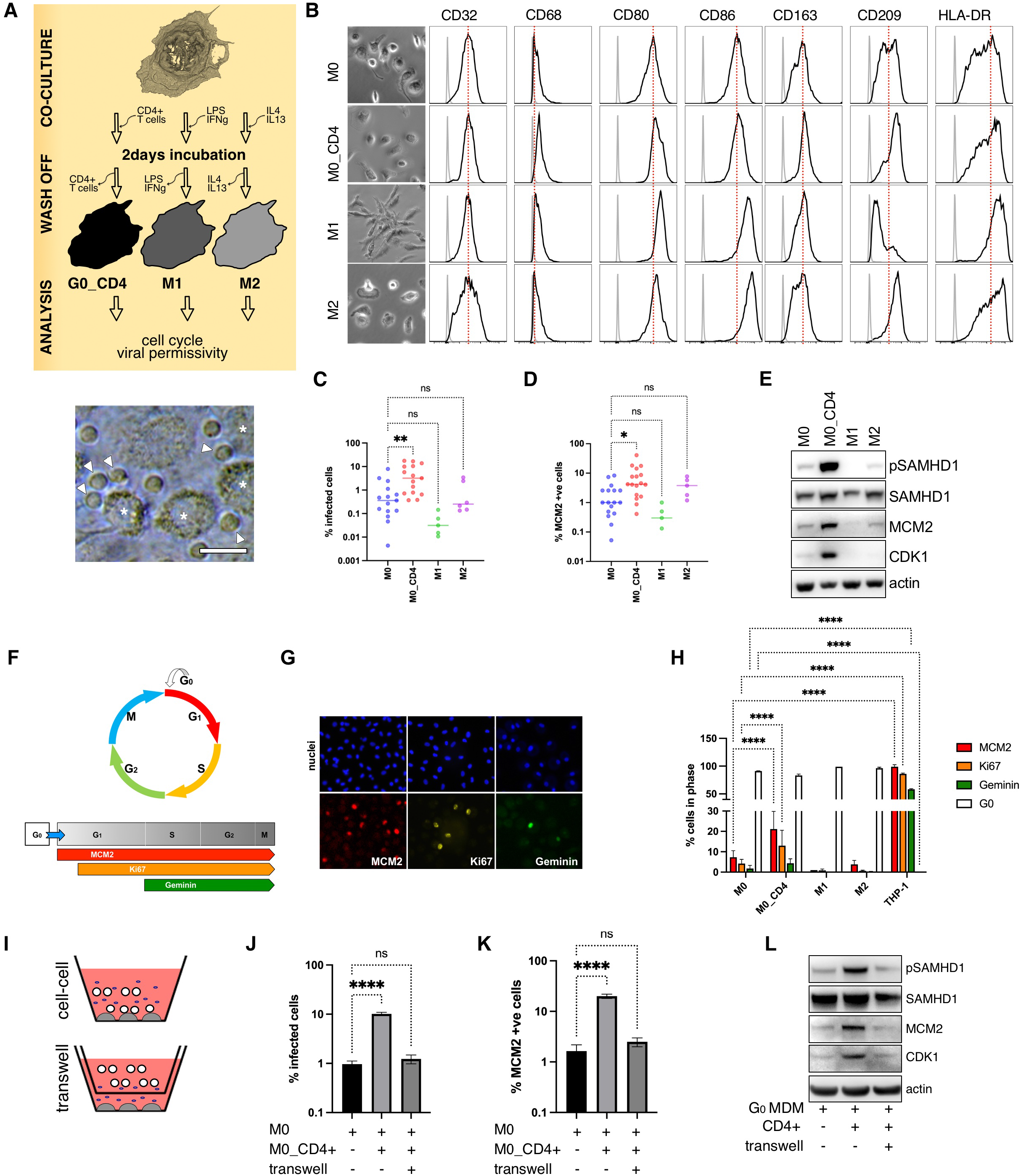
Direct contact with CD4+ T cells drives macrophage cell cycle re-entry with deactivation of SAMHD1 antiretroviral activity. **A**. Experimental design. Differentiated macrophages were co-cultured with activated autologous CD4+ T-cells, or stimulated by LPS/Interferon gamma (to polarize cells into M1 macrophages), or IL-4/IL-13 (into M2 macrophages). CD4+ T-cells were washed off 2 days later and cell cycle, viral permissivity was analysed. A representative example of microscopic field. * macrophages; scale bar:20um. **B**. Cell surface markers of macrophage polarization. Macrophages were stained for 6 different markers of polarization and analysed by flow cytometry. M0 (un-treated macrophages) are non-stimulated M-CSF differentiated cells; M0_CD4 (M0 macrophages after co-culture with T-cells); M1 (LPS/Interferon gamma) and M2 (IL-4/IL-13) macrophages. **C**. Macrophages were infected with VSV-G pseudotyped HIV-1 expressing GFP. Cells were fixed 2 days post-infection and percentage of GFP positive cells was quantified by using automated microscopic platform. Graphs represent average of n=15 (M0, M0_CD4); n=6 (M1, M2) biological replicates. Statistical analysis was performed using one-way ANOVA with Dunnett’s multiple comparisons test. ns, non-significant; *p < 0.1; **p < 0.01. Bars indicate mean with SD. **D**. Macrophages were fixed and stained for MCM2 (a marker of cell cycle re-entry) and percentage of positive cells was determined by using automated microscopic platform. Graphs represent average of n=16 (M0, M0_CD4); n=4 (M1, M2) biological replicates. Statistical analysis was performed using one-way ANOVA with Dunnett’s multiple comparisons test. ns, non-significant; *p < 0.1; **p < 0.01. Bars indicate mean with SD. **E**. Macrophages were lysed and cell lysates from a representative donor were used for immunoblotting. CDK1, MCM2 and SAMHD1 phosphorylation are markers of cell-cycle re-entry. **F**. A diagram of cell cycle depicting an expression of several cell cycle markers. None of the markers are present in G0; MCM2 is expressed in G1, S, G2,M; Ki67 is expressed in late G1, S, G2,M; geminin is expressed in S, G2,M. **G-H**. Macrophages were fixed and stained for cell cycle markers. **(G)** A representative acquisition field from macroscopic platform. **(H)** Percentage of positive cells was quantified by automated macroscopic platform and ImageJ. At least 10e4 cells was aquired and used for analysis. n=3 biological replicates; Ordinary two-way ANOVA with Sidak’s multiple comparisons test: ****p < 0.0001. Bars indicate mean with SD. **I-K**. Macrophages were plated at the bottom of transwell, CD4+ T-cells were added directly to macrophages to allow direct cell-cell contact or added in the top of transwell to prevent direct contact. **(J**,**K)** CD4+ T-cell were washed off 2 days later, **(J)** infected (HIV-1 GFP) or stained **(K)** for MCM2. Percentage of GFP or MCM2 positive cells was quantified by using automated microscopic platform. n=3 biological replicates. Statistical analysis was performed using one-way ANOVA with Dunnett’s multiple comparisons test. ns, non-significant; ****p < 0.0001. Bars indicate mean with SD. **L**. Macrophages were lysed and cell lysates from a representative donor were used for immunoblotting. CDK1, MCM2 and SAMHD1 phosphorylation are markers of cell-cycle re-entry.

Macrophages co-cultured with CD4 T cells were an order of magnitude more susceptible to infection compared to M0 macrophages (Figure 1C), co-incident with increased expression of intracellular MCM2 (Figure 1D) in co-cultured macrophages. Western blot analysis confirmed higher cellular expression of MCM2 under co-culture conditions and also demonstrated increased CDK1 expression and SAMHD1 phosphorylation with little change in overall SAMDH1 expression (Figure 1E), changes that are associated with cell cycle progression in MDM^9^.

We next performed more extensive analysis of cell cycle status to quantify cell cycle progression markers Geminin, MCM2 and Ki67 (Figure 1F,G). In this analysis, THP-1 monocytic tumour cells demonstrated high expression of all 3 markers, consistent with actively dividing cells. Co-cultured macrophages demonstrated significantly higher expression of MCM2 and Ki67 compared to M0 macrophages, indicating G0 to G1 progression. The modest elevation in Geminin expression was not statistically significant, supporting minimal progression to S, and G2/M.

We next sought to establish whether direct cell-cell contact was required for CD4 T cells to induce cell cycle progression in macrophages. Using a transwell system (Figure 1I) we observed that placement of a barrier between macrophages and T cells abrogated the increased susceptibility to infection with PV particles, as well as the cell cycle progression (Figure 1J-K). Immunoblot confirmed that the transwell prevented an increase in MCM2, CDK1 and pSAMHD1 protein levels following the addition of CD4 T cells (Figure 1L). We conclude that direct cell contact is required for T cells to induce cell cycle progression and relieve viral restriction in macrophages.

In order to elucidate pathways that might be involved in mediating CD4 T cell contact dependent cell cycle progression in macrophages we performed phosphoproteomic analysis. Three time points were assessed: 0, 5 and 120 minutes following contact between cells and cell cycle progression in macrophages was separately confirmed by immunoblot at 2 days timepoint (Figure 2A-B). A number of phosphoproteins were either upregulated or downregulated as demonstrated by volcano plots with the largest number of proteins changing between 0 and 120 minutes (Figure 2C). Gene Ontology and Ingenuity pathway analysis showed that the frequently altered phosphoproteins were involved in signal transduction pathways (Figure 2D). Given this result, activation of signal transduction and our previous work recognising the importance of the MEK/ERK pathway in the activation of macrophage G0-G1 transitioning^9^, we hypothesised that MEK/ERK pathway is involved in CD4 T cell contact dependent cell cycle progression in macrophages. To test whether this pathway was responsible for the cell contact phenotype we used inhibitors of MEK (U0126), CDK4/6 (Palcociclib) and CDK1 (RO-3306). Each of the inhibitors almost completely inhibited MCM2, CDK1 and pSAMHD1 expression following cell-cell contact, indicative of cell cycle arrest in G0 (Figure 2E). This was reflected in concomitant blockade of infection with HIV-1 PV following drug treatment (Figure 2F).

**Figure 2:**
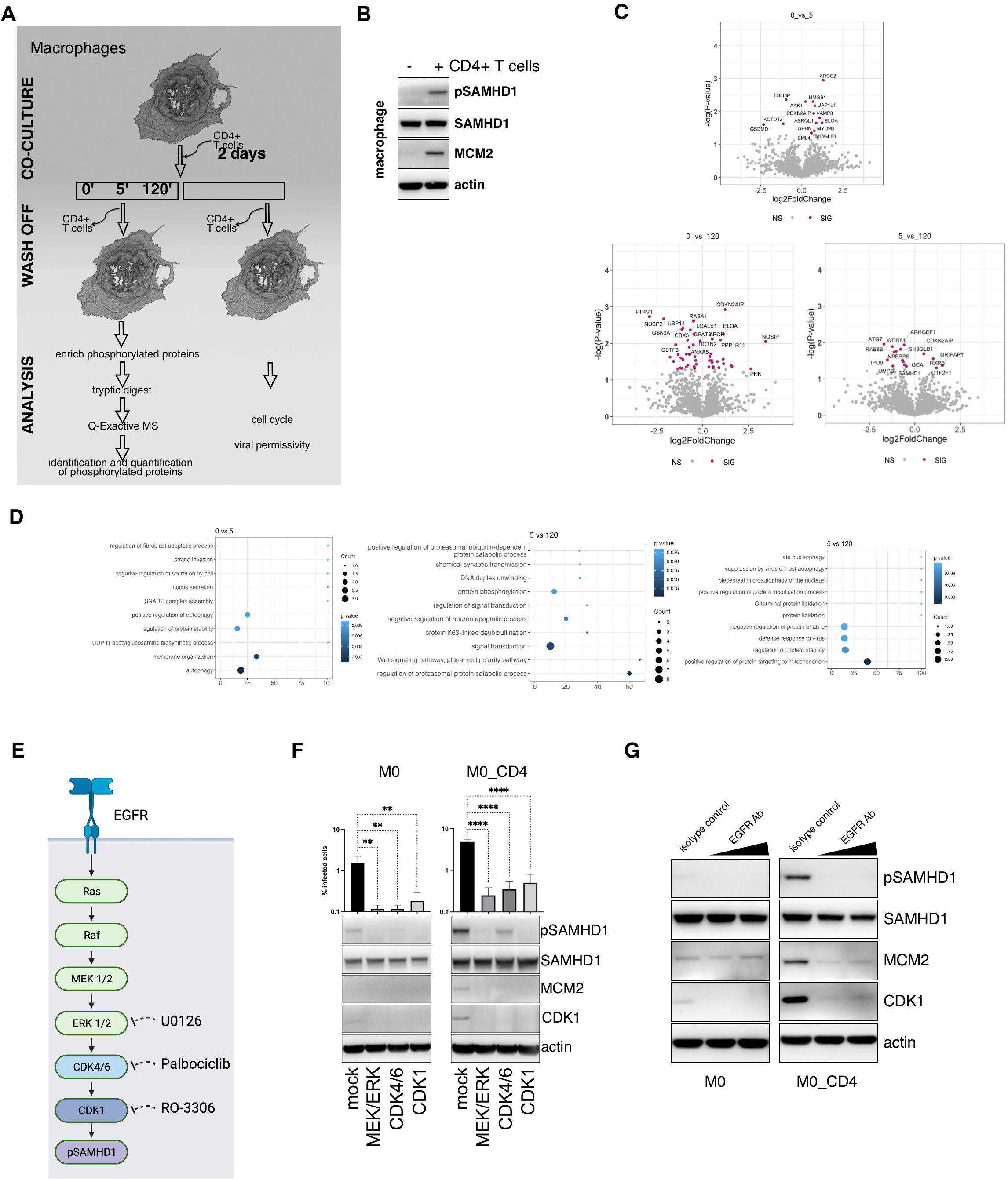
Direct cell-to-cell contact drives cell cycle re-entry in human macrophages through EGF receptor and canonical MEK/ERK signalling. **A**. Experimental design. Differentiated macrophages were co-cultured with activated autologous CD4+ T-cells for 0min, 5min, 120min. T-cells were washed off and macrophages lysed. Proteins were extracted and phosphoproteins enrished. Proteaome and Phosphoproteome were performed using 2 donors. **B**. Macrophages were co-cultured with CD4+ T-cells and T-cells were washed off 2 days later. Macrophages were lysed and cell lysates from one of the donors were used for immunoblotting. CDK1, MCM2 and SAMHD1 phosphorylation are markers of cell-cycle re-entry. **C**. Volcano plot showing differentially expressed phosphoproteins in macrophages following CD4+ T cell contact versus the absence of T cell contact. Coloured proteins were significantly up or down regulated with significance level of 0.1. **D**. Number of proteins observed to be differentialy phosphorylated. Analysis comparing phosphoproteins from 0 vs. 5min; 5 vs. 120min; and 0 vs. 120min. **E**. Protein ontology. Identified changes to protein phosphorylation were classified accoding to protein biological functions. **F**. Diagram of canonical MAPK pathway with highligted inhibitors of kinases. **G**. Macrophages were untreated (M0) or co-cultured with CD4+ T-cells for 2 days in the presence or absence (mock) of inhibitors. T-cells were washed off and macrophages infected with VSV-G pseudotyped HIV-1 expressing GFP. Percentage of GFP positive cells was quantified by using automated microscopic platform 2 days later. Macrophages were lysed and cell lysates from a representative donor were used for immunoblotting. CDK1, MCM2 and SAMHD1 phosphorylation are markers of cell-cycle re-entry. n=3 biological replicates. Statistical analysis was performed using one-way ANOVA with Dunnett’s multiple comparisons test. **p < 0.01; ****p < 0.0001. Bars indicate mean with SD. **H**. Macrophages were co-cultured with CD4+ T-cells for indicated time. T-cells were washed off and macrophages lysed. Cell lysates from a representative donor were used for immunoblotting. **I**. Macrophages were untreated (M0) or co-cultured with CD4+ T-cells for 2 days in the presence of control isotype antibody or increasing concentration of anti-EGFR antibody. T-cells were washed off and macrophages lysed. Cell lysates from a representative donor were used for immunoblotting.

Growth factors and mitogens use the canonical Ras-Raf/MEK/ERK signalling cascade to transmit signals from their receptors to regulate gene expression^16^. EGFR is one of the most important receptors in this pathway^17^. We sought to test whether this receptor could be involved in initiating macrophage cell cycle progression. We therefore treated macrophages with a monoclonal antibody to EGFR prior to co-culture with CD4 T cells. The antibody prevented cell cycle progression even in the presence of CD4 T cells based on the expression of cell cycle associated markers MCM2 and CDK1, and pSAMHD1 (Figure 2E,G). This suggests that EGFR signalling may contribute to cell cycle state in macrophages, and that it can be activated to induce potent cell cycle progression following CD4 T cell contact.

## Discussion

These experiments demonstrating cell cycle progression following antigen independent T cell contact delineate a new phenotype of macrophage, with distinct distribution of cell surface marker distribution from M0/M1/M2. Phosphoproteomic analysis revealed that within hours of contact, cell cycle associated proteins in the Ras-Raf-MEK-ERK axis were activated, and this was confirmed biochemically through inhibition of this pathway using immunoblot.

EGFR appeared to mediate the phenotype, consistent with cell-cell signalling. The cell cycle cascade in macrophages resulted in G0-G1 transition as demonstrated by nuclear staining for Ki67 and MCM2, and deactivation of the antiviral activity of SAMHD1. Tissues such as lymph nodes are characterised by close cell contact between T cells and macrophages^15^. The deactivation of SAMHD1 antiviral activity following cell contact may underlie observations that macrophages are a reservoir for HIV^18,19^ and lymph nodes remain a difficult to treat HIV reservoir even in the face of antiretroviral therapy^20^.

## Methods

### Cells, plasmids and viruses

293T (a human embryonic kidney cell line, ATCC CRL-3216) cells were maintained in Dulbecco’s modified Eagle medium (DMEM) supplemented with 10% foetal calf serum (FCS), 100 U ml^−1^ penicillin and 100 mg ml^−1^ streptomycin, and regularly tested and found to be mycoplasma free. THP-1 cells, a kind gift from G. Towers, maintained in RPMI (DMEM) supplemented with 10% FCS, 100 U ml^−1^ penicillin and 100 mg ml^−1^ streptomycin. pBOB-EF1-FastFUCCI-Puro was a gift from Kevin Brindle & Duncan Jodrell (Addgene plasmid # 86849; http://n2t.net/addgene:86849; RRID: Addgene_86849) (Koh *et al*, 2017).

#### Reagents, inhibitors, antibodies

All chemicals were purchased from Sigma unless indicated otherwise. Kinase inhibitors used: CDK4/6 inhibitor (PD 0332991, Palbociclib) from Sigma; MEK/ERK inhibitor U0126 from Calbiochem (San Diego, USA); Antibodies used were as follows: anti-cdc2 (Cell Signaling Technology, Beverly, MA, USA); anti-SAMHD1 (ab67820, Abcam, UK), beta-actin (ab6276, abcam, UK); mouse anti-MCM2 (BM-28, BD Biosciences, UK); and rabbit anti-MCM2 (SP85) from Sigma; pSAMHD1 ProSci (Poway, CA, USA); anti-Geminin (NCL-L-Geminin, Leica); anti-Ki67 (ab15580, abcam); anti-human CD32, CD68, CD163, CD80, CD86, CD209, HLA-DR (BD, 550586, 562117, 563887, 340294, 555666, 551545, 567054). Kinase inhibitors used are CDK4/6 inhibitor (PD 0332991, Palbociclib, Sigma); MEK/ERK inhibitor U0126 (Calbiochem); CDK1 inhibitor (RO-3306, Sigma).

### Monocyte isolation and differentiation and CD4 isolation

PBMC were prepared from apheresis cones from NHS Blood Center Cambridge by density-gradient centrifugation (Lymphoprep, Axis-Shield, UK). MDM were prepared by adherence with washing of non-adherent cells after 2 h, with subsequent maintenance of adherent cells in RPMI 1640 medium supplemented with 10% human serum or 10% foetal calf serum and MCSF (10 ng/ml) for 3 days and then differentiated for a further 4 days in RPMI 1640 medium supplemented with 10% human/foetal calf sera without M-CSF. Human AB serum (Sigma) was used to prepare unstimulated cells or FCS (Biosera or Sigma) to prepare stimulated cells. Autologous CD4^+^ T cells were isolated from PBMCs with a negative CD4^+^ T cell isolation kit (130-096-533, Miltenyi Biotec).

### Cell cycle analysis using fluorescence ubiquitination cell cycle indicator (Fucci)

Fucci cassette was cloned from pBOB-EF1-FastFucci-Puro vector to pEXN-MNCX using BamHI/NotI restriction sites. Fucci-containing lentiviral particles were produced as follows: 293Tv cells were transfected with pEXN-MNCX-Fucci, CMVi and pMD2.G. Cell supernatants containing viruses (Fucci VLP) were collected 48 h post-transfection and frozen at −80°C. Macrophages were transduced using Fucci VLP for 18 h.

### SDS PAGE and immunoblot

Cells were lysed in reducing Laemmli SDS sample buffer containing PhosSTOP (Phosphatase Inhibitor Cocktail Tablets, Roche, Switzerland) at 96°C for 10 min and the proteins separated on NuPAGE^®^ Novex^®^ 4–12% Bis–Tris Gels. Subsequently, the proteins were transferred onto PVDF membranes (Millipore, Billerica, MA, USA), the membranes were quenched and proteins were detected using specific antibodies. Labelled protein bands were detected using Amersham ECL Prime Western Blotting Detection Reagent (GE Healthcare, USA) and ChemiDoc MP Imaging System (Bio-Rad) CCD camera. Protein band intensities were quantified using ChemiDoc MP Imaging System and Image Lab software (Bio-Rad, Hercules, CA, USA).

### Immunofluorescence

Cells were fixed in 4% PFA, quenched with 50 mM NH_4_Cl and permeabilised with 0.1% Triton X-100 in PBS. After blocking in PBS/1% FCS, cells were labelled for 1 h with primary antibodies diluted in PBS/1% FCS, washed and labelled again with Alexa Fluor secondary antibodies for 1 h. Cells were washed in PBS/1% FCS and stained with DAPI in PBS for 5 min. Labelled cells were detected using ArrayScan high-content system (Thermo Fisher, Waltham, MA, USA) and analysed using Harmony (PerkinElmer, Waltham, MA, USA) and ImageJ software.

### VSV-G pseudotyped virus single-round infection

VSV-G-pseudotyped HIV-1 GFP expressing viruses were added to MDM. 4h post-incubation, the inoculum was removed and cells were washed once in a culture medium. This was left for 36 hr post infection before the cells were stained by Hoechst for nuclei. The percentage of infected GFP-expressing cells versus total cells was determined 48 h post-infection using ArrayScan high-content system (Thermo Fisher, Waltham, MA, USA) and analysed using Harmony (PerkinElmer, Waltham, MA, USA) and ImageJ software.

### Proteomics

Phosphoproteins of donor cells were enriched with the Phosphoprotein Enrichment Kit (Thermo Pierce) according to manufacturer’s instructions and subsequently prepared in parallel for analysis by LC-MS/MS in parallel with not enriched lysates.

Briefly, samples were reduced with DTT (5mM, 30 minutes at room temperature), alkylated with Iodoacetamide (20mM, 30 minutes in the dark), subjected to protein precipitation using Chloroform/Methanol and digested with trypsin (Promega). Desalted and acidified peptides (0.1% TFA, SOLA (Thermo Pierce)) were injected into a LC-MS/MS platform consisting of Q-Exactive mass spectrometer and Dionex Ultimate 3000 UPLC, after separation on a EASY Spray column (50cm, all Thermo) with a gradient of 2-35% acetonitrile in 0.1% formic acid/5% DMSO over 60 minutes. Data were acquired in DDA mode using standard parameters and imported into Progenesis QI (Waters) for label free quantitation (identification through Mascot 2.5 (Matrix Science)). Peptides failing the 1% FDR threshold and with a Mascot score <20 were excluded from analysis. The mass spectrometry proteomics data have been deposited to the ProteomeXchange Consortium via the PRIDE partner repository (https://www.ebi.ac.uk/pride/) with the dataset identifier PXD048462 and 10.6019/PXD048462.

## Notes

### Competing Interest Statement

The authors have declared no competing interest.

### Summary of Updates

references added and abstract amended

